# Age dictates brain functional connectivity and axonal integrity following repetitive mild traumatic brain injuries

**DOI:** 10.1101/2024.01.25.577316

**Authors:** Marangelie Criado-Marrero, Sakthivel Ravi, Ekta Bhaskar, Daylin Barroso, Michael A. Pizzi, Lakiesha Williams, Cheryl L. Wellington, Marcelo Febo, Jose Francisco Abisambra

**Author notes:** These authors contributed equally to this work (Marangelie Criado-Marrero, Sakthivel Ravi).

## Abstract

Traumatic brain injuries (TBI) present a major public health challenge, demanding an in-depth understanding of age-specific signs and vulnerabilities. Aging not only significantly influences brain function and plasticity but also elevates the risk of hospitalizations and death following repetitive mild traumatic brain injuries (rmTBIs). In this study, we investigate the impact of age on brain network changes and white matter properties following rmTBI employing a multi-modal approach that integrates resting-state functional magnetic resonance imaging (rsfMRI), graph theory analysis, diffusion tensor imaging (DTI), and Neurite Orientation Dispersion and Density Imaging (NODDI). Utilizing the CHIMERA model, we conducted rmTBIs or sham (control) procedures on young (2.5-3 months old) and aged (22-month-old) male and female mice to model high risk groups. Functional and structural imaging unveiled age-related reductions in communication efficiency between brain regions, while injuries induced opposing effects on the small-world index across age groups, influencing network segregation. Functional connectivity analysis also identified alterations in 79 out of 148 brain regions by age, treatment (sham vs. rmTBI), or their interaction. Injuries exerted pronounced effects on sensory integration areas, including insular and motor cortices. Age-related disruptions in white matter integrity were observed, indicating alterations in various diffusion directions (mean, radial, axial diffusivity, fractional anisotropy) and density neurite properties (dispersion index, intracellular and isotropic volume fraction). Inflammation, assessed through Iba-1 and GFAP markers, correlated with higher dispersion in the optic tract, suggesting a neuroinflammatory response in aged animals. These findings provide a comprehensive understanding of the intricate interplay between age, injuries, and brain connectivity, shedding light on the long-term consequences of rmTBIs.

## Introduction

Traumatic brain injuries (TBI) profoundly impact the global population affecting lifestyle and work productivity of an estimated 50 million individuals worldwide^1^. They are a leading cause of disability imposing high economic burden of health costs for the individual and healthcare systems^2,3^. Among TBIs, repeated mild traumatic brain injuries (rmTBIs), commonly recognized as sequential and multiple concussions, are prevalent and underdiagnosed traumas^3^. Given the mild nature of these injuries and cofounding symptoms with other diseases, signs may not be noticeable shortly after the incident, which presents substantial challenges to differentiate temporary effects and identify profound pathogenic signatures in brain^3,4^.

Repetitive mTBIs are a major environmental risk factor for developing post-traumatic stress disorder (PTSD) and neurodegenerative conditions such as Alzheimer’s disease and related dementias (ADRD)^5–8^. Aging is a significant factor contributing to the elevated risk of these disorders and the occurrence of traumatic brain injury-related cases and fatalities^9,10^. However, whether aging and rmTBI, alone or in combination amplify brain dysfunction remains largely undefined. Identifying the intricate relationship between aging and rmTBI on neurodegenerative conditions is critical and urgent.

Resting-state functional magnetic resonance imaging (rsfMRI) and diffusion tensor imaging (DTI) are valuable non-invasive tools used to define imaging biomarkers of brain damage after injury^11–17^. rsfMRI enables the evaluation of functional connectivity, whereas DTI captures structural alterations in the white matter fiber tracks. Combined with graph theory, these imaging techniques determine changes in topological brain networks and subnetworks, functional connections between regions of interest, morphology following rmTBI^15,18–20^, and rmTBI associations with cognitive decline and neurodegeneration^17,21^.

Young and aged adults show a higher prevalence for TBI cases and deaths, as reported by Center of Disease Control (CDC)^3^. Therefore, we used rsfMRI and graph theory to delineate distinct connectivity patterns among 148 regions in the brain of young (2.5-3 months old) and aged (22 months old) mice to model the higher-risk age groups in humans. Additionally, we assessed white matter integrity and its compartments by employing DTI and neurite orientation dispersion and density imaging (NODDI) techniques. We identified significant alterations in functional connectivity (79 regions) and axonal diffusivity (16 regions) affected by age and treatment (sham vs. rmTBI). Notably, age-related disparities in network connectivity were markedly found in eigenvector centrality, which quantifies the importance of connections between the most influential regions in the brain network. This study represents an important step in characterizing age-related differences in connectivity during the initial days after rmTBI. These results provide a baseline for comprehensive network effects subject to age and injury. Long-term, we aim to translate these results for more effective diagnosis, brain activity monitoring, recovery prediction, and assessment of elderly patients with rmTBI.

## Methods

### Animal husbandry

We obtained C57BL/6J wild-type young mice of both sexes from Jackson Laboratory (Bar Harbor, ME, USA). The aged wild-type mice were received from Dr. Dave Borchelt, University of Florida. Animals were housed and maintained under a 12-hour light cycle in standard room conditions with free access to water and food. All animal procedures were approved by the University of Florida’s Institutional Animal Care and Use Committee (IACUC) and adhered to the guidelines outlined in the National Institutes of Health Guide for the Care and Use of Laboratory Animals.

### Closed-Head Impact Model of Engineered Rotational Acceleration (CHIMERA)

We used CHIMERA to induce rmTBI following previously published protocols^17,22^. Male (n = 23) and female (n = 10) mice were randomly assigned to either sham or CHIMERA injury procedures at their respective age groups: ‘young’ at 2.5-3 months and ‘aged’ at 22 months old. Mice received isoflurane (maintained at 1-2.5%) during the injury or sham protocols. The injured group received two mild closed head impacts (0.6 J) 24 hours apart; the sham group did not sustain impacts. Mice were injected subcutaneously with meloxicam (10 mg/kg) to control pain within 48 hours. Tissue collection was performed at either 7 days (young) or 26 days (aged) after injury (**Figure 1A**). This study only includes tissue examination in the aged group. Immunohistochemistry results for gliosis and other disease-like markers in young mice were reported previously^17^.

**Figure 1.**
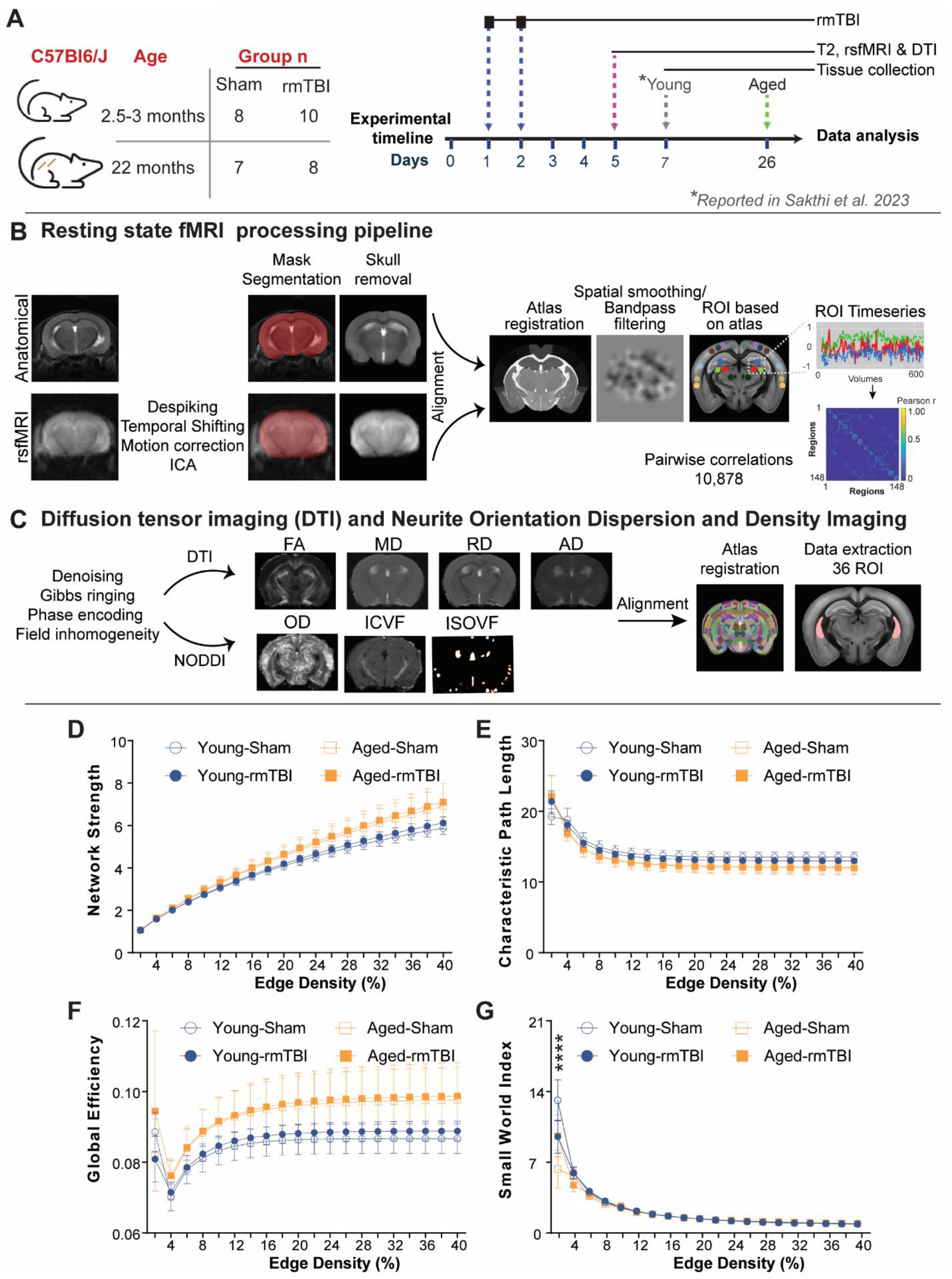
Experimental Design and Neuroimaging Analysis. (**A**) Schematic representation showing the number of rmTBIs, rsfMRI and tissue collection. (**B**) Resting state functional MRI processing pipeline. (**C**) Diffusion tensor imaging (DTI) and neurite orientation dispersion and density imaging (NODDI) workflow. Brain network was assessed by using (**D**) network strength (measures overall network connectivity), (**E**) characteristic path length (quantifies network communication efficiency), (**F**) global efficiency (assesses information transmission efficiency), and (**G**) small-world index (evaluates local and global information balance). Repeated measures two-way ANOVA was conducted in all networks analyses examining a range from 2 to 40 edge density thresholds. TBI, trauma brain injury; ROI, region of interest; rsfMRI, resting state functional magnetic resonance imaging; ICA, independent component analysis; FA, fractional anisotropy; MD, mean diffusivity; AD, axial diffusivity; RD, radial diffusivity; ISOVF, isotropic volume fraction; ICVF, intracellular volume fraction; OD, orientation dispersion index.

### Imaging procedures

#### Image collection

Five days post-injury (dpi), anatomical (T2), diffusion-weighted (dwi) and functional (fMRI) images were collected from animals anesthetized with isoflurane (4% for initiation and 1% during maintenance) and medical grade air (70% N_2_: 30% O_2_). Image scanning was completed at the Advanced Magnetic Resonance Imaging and Spectroscopy Facility at the University of Florida in Gainesville, FL using a 11.1T MRI scanner (Magnex Scientific Ltd, Oxford, UK). Each animal was positioned to minimize motion artifacts under a radiofrequency (RF) coil tuned to 470.7MHz (1 H resonance). This coil generated electromagnetic fields to detect nuclear magnetic resonance signals. A T2-weighted TurboRARE sequence was used with parameters: TE (echo time) = 41.42 msec, TR (repetition time) = 4000 msec, RARE factor = 16, average number = 12, FOV (field of view) 15 mm × 15 mm, 0.9 mm slice thickness, and a data matrix of 256 × 256 with 14 interleaved ascending coronal slices. Resting-state fMRI were acquired via a single-shot spin echo (SE) echo planar image (EPI) blip-up and blip-down sequence with parameters: TE = 15 msec, TR = 2000 msec, 600 repetitions, and a data matrix of 64 × 48 with 14 interleaved ascending coronal slices in the same space as the reference scan (anatomical).

For diffusion, image acquisition was completed in four shot 2-shell SE-EPI using the following parameters: TE = 19 msec, TR = 4000 msec, average number = 4, gradient duration = 3 msec, diffusion time spacing = 8 msec, FOV = 11 mm x 0.11 mm, slice thickness = 0.7 mm, matrix = 128 x 96, slice per mouse = 17. For DTI analysis, 47 images were collected, a B0 baseline image and 46 images with gradient directions for b values of 600 and 2000 sec/mm^2^.

#### Image preparation

The University of Florida HiPerGator supercomputer was used to complete all image processing. Images were converted from Bruker to nifty format. All rsfMRI images underwent a comprehensive pre-processing pipeline using Analysis of Functional NeuroImages (AFNI)^23^, Functional MRI of the Brain Library (FSL)^24^, Advanced Normalization Tools (ANTs) software as outlined previously^17^. Pre-processing involved distortion and phase correction using FSL’s TOPUP, removal of time series spikes and slow temporal variations, noise decomposition applying the independent component analysis (ICA), removal of skull using three-dimensional pulsed coupled neural networks (PCNN3D) in MATLAB (**Figure 1B**). Manual correction of image masks was performed using ITK-SNAP^25^. Additional processing steps involved spatial smoothing (0.4 mm), and extraction of time series data. Time series from functional images, divided into 600 volumes, were linearly aligned (using FSL FLIRT) and nonlinearly transformed (using ANTS) to the anatomical image. Then, these were aligned and registered to the two times down sampled parcellated mouse common coordinate framework (version 3, or CCFv3)^26^ which was used as template to determine coordinates corresponding to 148 regions of interest (ROI) across the brain (**Supplemental Table 1**).

#### Network metrics in rsfMRI

Timeseries correlations among ROIs were calculated by Pearson’s coefficient resulting in 10,878 relationships per brain. Fisher transformation was applied to normalize correlation coefficients before running analyses. The Brain Connectivity toolbox in MATLAB identified complex network properties and regional interactions. Network analysis was calculated using established methodologies^17,27^. Edge density thresholds ranging from 2% to 40% were systematically applied to evaluate a spectrum of global network metrics capturing strongest connections. Network strength measures the overall connectivity computed as the sum of edge weights connected to a region (node) indicating strength of communication between these connections. The characteristic path length measures the efficiency of information transfer, capturing how readily information spreads across the network. Global efficiency quantifies the brain’s overall ability to integrate information across distant regions. The Small world index (SWI) represents a balance between local clustering and global efficiency, offering a more detailed understanding of network topology. For ROI-specific measures, a uniform 10% threshold was used as this edge density may provide a comprehensive view of both strong and moderate connections in the brain network. BrainNet Viewer was used to illustrate brain connectome maps^28^.

#### Diffusion tensor imaging (DTI) and Neurite Orientation Dispersion and Density Imaging (NODDI)

Bruker diffusion weighted images were converted to nifty files and then small and large shells were combined into one image using fslmerge. Images with severe noise (exhibiting high interference, random fluctuations in the signal, abnormal motion distortions) were excluded from the study. Remaining images were converted to mif file for Mrtrix software processing. Denoising (enhancing signal-to-noise ratio), Gibbs ringing (suppressing artifactual oscillations), phase encoding (correcting for motion distortions), B1 field inhomogeneity (mitigating variations in the RF field) were applied to the data (**Figure 1C**). These steps ensure higher accuracy and uniform signals across the brain before analysis.

For DTI, FSL’s dtifit and CAMINO were used for linear and nonlinear diffusion tensor model application into our diffusion-weighted data computing fractional anisotropy (FA), mean diffusivity (MD), radial diffusivity (RD), and axial diffusivity (AD) value in square millimeters per second (mm^2^/s). For NODDI, 54 volumes composed of B0 (total of 2), b value of 60 sec/mm^2^ (total of 6), and b value of 2000 sec/mm2 (total of 46) images were processed adapting published protocols^29^. To characterize microstructural properties, the orientation dispersion (OD), intra-cellular volume fraction (ICVF), and isotropic volume fraction (ISOVF) was calculated in neural tissues using NODDI AMICO software package^30^. Preprocessed DTI and NODDI images were then aligned to the mouse CCFv3 reference atlas. We first applied linear and nonlinear registrations between FA images, which has the best resolution among the DTI scans, to the reference atlas. The resulting linear transformations in FA were applied to the MD, RD, AD, OD, ISOVF, and ICVF images. After all images were aligned to a standard space, a common mask was applied to exclude any extra borderlines in images. Guided by atlas parcellations, manual segmentations were drawn for 36 ROIs in a common group template using ITK-SNAP (**Supplemental Table 2**). A single value corresponding to mean voxel intensity was extracted per region of interest (ROI) using the FSL fslstats function.

### Immunohistochemical analysis

Ionized calcium-binding adapter molecule 1 (Iba-1) and glial fibrillary acidic protein (GFAP) staining was performed as previously established^17^. In brief, 5 µm coronal brain sections underwent deparaffinization, rehydration, and antigen retrieval. Endogenous peroxidase was eliminated using the mixture of 0.3% H_2_O_2_ and 10% Triton X-100, and sections were blocked with 10% goat serum. Tissue was incubated overnight with anti-Iba-1 (1:1000, PA5-27436, Invitrogen) and anti-GFAP (GA5) (1:1000, CS3670S, Cell Signaling) antibodies followed by 30 min incubation with biotinylated secondary antibodies. Sections were then treated with an avidin-biotin complex (ABC) reagent and developed in DAB (Cat# 5510-0031). Image quantification used Image Scope software (v12.4.3.5008) with positive pixel count program. Average values from three sections per brain determined Iba-1 and GFAP expression.

### Statistical analysis

Network and tissue analyses were conducted using GraphPad Prism software (version 10.0.3). Global network outcomes were described using repeated measures two-way ANOVA, correcting for multiple comparisons with the Tukey test. All statistical analyses for the imaging data were performed using R (v4.1.2). Before running analyses, data were assessed for normal distribution and homogeneity of variances. Extreme outliers were identified using the interquartile range (IQR) method. Main effects of age (young vs. aged) and treatment (sham vs. rmTBI) were tested by type III two-way ANOVA on each ROI with false discovery rate Benjamini-Hochberg (5% FDR-BH) correction. ROIs resulting in statistically significant (p value < 0.05) by main effects were followed by post-hoc analyses. Significant differences between group pairs were accounted when pairwise comparisons with FDR-BH shown p < 0.05 together with a large Cohen’s effect size (+/− 0.8) between comparing means. Two-tail unpaired Welch’s t-test were used for glial analyses.

## Results

### Experimental design and global network properties

Aging significantly affects brain function and plasticity after injuries^29,31–34^. To investigate the extent to which age and rmTBIs influence overall brain network stability and white matter properties, we performed CHIMERA injuries or sham control procedures (2 injuries, 24 hours apart, 0.6 J) on 2.5-3 and 22 months of age male and female mice. Image scanning of rsfMRI and DTI was done 5 days post-injury (dpi) (**Figure 1A-1C**). In functional images, we applied graph theory matrices to assess overall network integrity: network strength (indicating global connectivity, **Figure 1D**), characteristic path length (information travel distance, **Figure 1E**), global efficiency (overall communication, **Figure 1F**) and small-world index (SWI; balancing local and global connectivity, **Figure 1G**). In general, major global differences were not detected except for a significant decreased in SWI induced by aging, indicating reduced communication efficiency between various brain regions. However, injuries induced an opposing effect on SWI in different age groups. Specifically, rmTBI reduced network segregation in young mice while enhancing it in aged animals, thereby influencing the formation of connections with neighboring areas.

### Functional connectivity changes at local regions affected by age and treatment

We then inspected how 148 regions of interest (ROI, nodes) distributed throughout the brain were functionally connected or restructured by age or after injury. We found that 79 out of the 148 regions were altered by age (young vs. aged), txt (sham vs. rmTBI), or the interaction between these factors (**Figure 2**). Color-coded regions represent areas showing a statistically significant value (p < 0.05) and a large effect size (Cohen’s d equal or +/− 0.8). **Figure 2A** highlights regions with at least 2 significant changes and substantial effect among the metrics. The most prominent changes were observed in eigenvector centrality, a metric that characterizes how age and rmTBI impact the number of connections and their influence on neighboring regions and community hubs in specific areas of the brain. Age emerged as the primary influencing factor on eigenvector centrality in 64% of regions (**Figure 2B**). Not surprisingly, injuries exerted greater effects on areas critical for sensory integration and pain such as the entorhinal cortex and agranular insula (posterior and ventral areas). The total number of affected regions in left and right hemispheres was counted to detect any asymmetric measures (**Figure 2C**). Interestingly, while fewer regions were affected in the left hemisphere, it’s noteworthy that this side exhibited more pronounced connectivity changes when considering all metrics within these regions. These findings emphasize the broad interplay of age, injury, and brain connectivity, providing a deeper understanding of numerous brain connections affected by TBIs.

**Figure 2.**
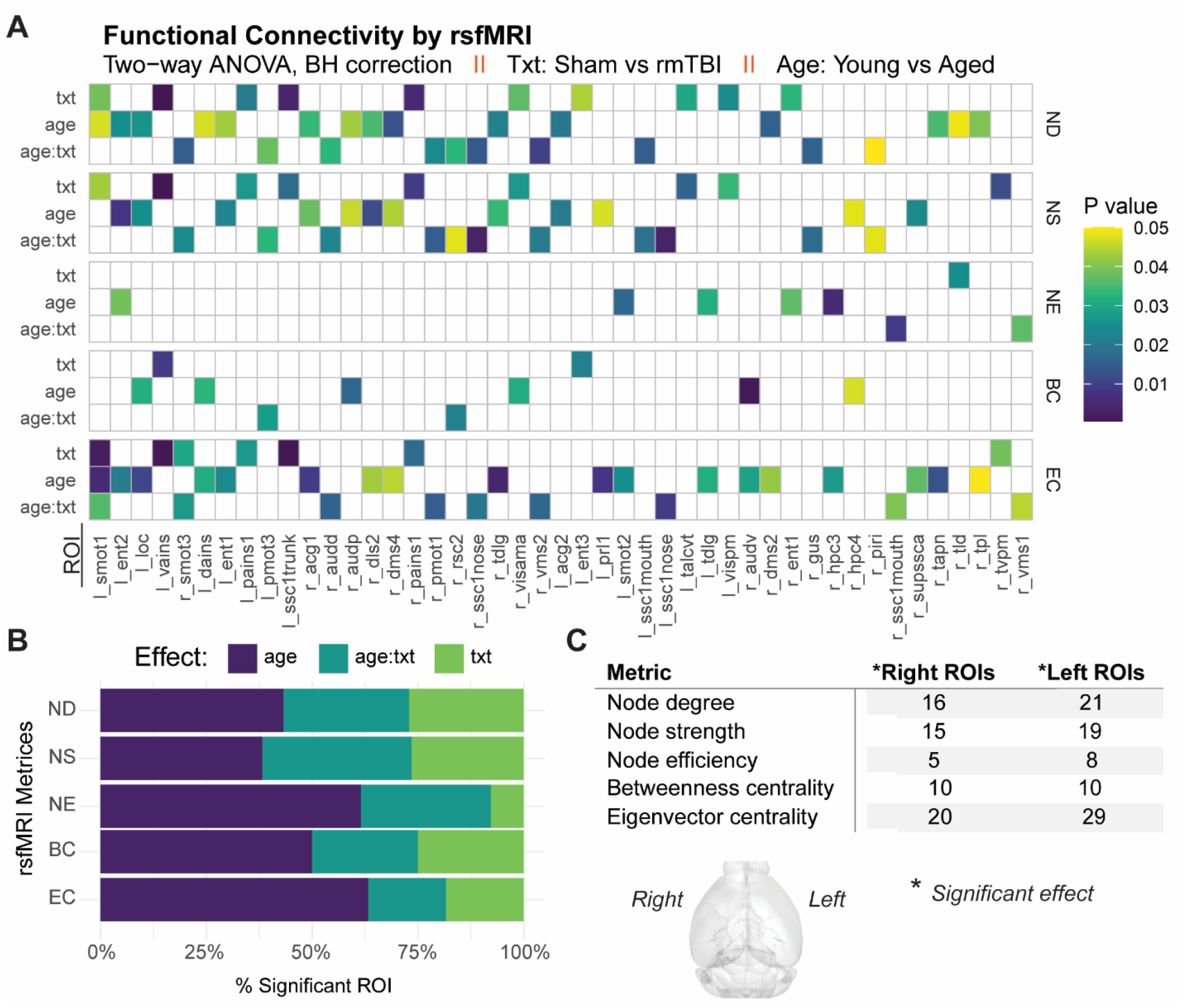
Age and Treatment Effects on rsfMRI Metrics in Brain. A threshold of 10% edge density was applied for local analyses, which is within the most robust and least dense connections. (**A**) Main effects and interaction of age and treatment (txt = sham vs rmTBI) on the following rsfMRI metrices: node degree (ND), node strength (NS), node efficiency (NE), betweenness centrality (BC), and eigenvector centrality (EC). Out of the 148 regions of interest evaluated, only those with 2 or more significant values are represented in the heatplot. Statistically significant (p < 0.05) effects for each region of interest (ROI) resulting from Type III two-way ANOVA with Benjamini false discovery rate (FDR-BH) correction are represented in color. White squares denote non-significant values. (**B**) Percentage of significant ROI affected by age, txt, interaction age and txt for each metric. (**C**) Significant main effects distribution in right and left-brain hemisphere. rsfMRI = resting state functional magnetic resonance imaging. ROI names are detailed in **Supplemental Table 1.**

### Age, brain network reorganization, and injury effects

To further investigate these effects, we performed multiple comparison test to pinpoint specific distinctions in brain regions among the groups (**Figure 3A**). This analysis revealed that the injuries primarily increased eigenvector centrality and node strength in motor cortices in young mice (**Figure 3B**). Conversely, in aged mice, injuries resulted in decline in both the number (degree) and strength of local connections within the brain. This suggests a diminished capacity for local interactions and communications. Additionally, these injuries affected the connections’ influence on neighboring areas, potentially disrupting the overall network dynamics and communication between different regions of the brain of aged mice. In young mice, the only region that exhibited a reduction in node strength, node degree and eigenvector centrality was the posterior agranular insula. In aged mice, the ventral segment of insula showed a decrease in the same measurements (**Figure 3C**). Minor differences between young and aged sham (Y-Sh vs A-Sh) were detected where aging heightened activity in auditory (betweenness and eigenvector centrality), hippocampal (node strength in CA3 area), while reducing number of connections in striatal areas. In the hippocampal region (dentate gyrus), aged-injured mice revealed increased influence and region strength of functional connections relative to young-rmTBI mice. These observations confirm that the effect of repetitive brain injuries on brain connectivity varies significantly with age.

**Figure 3.**
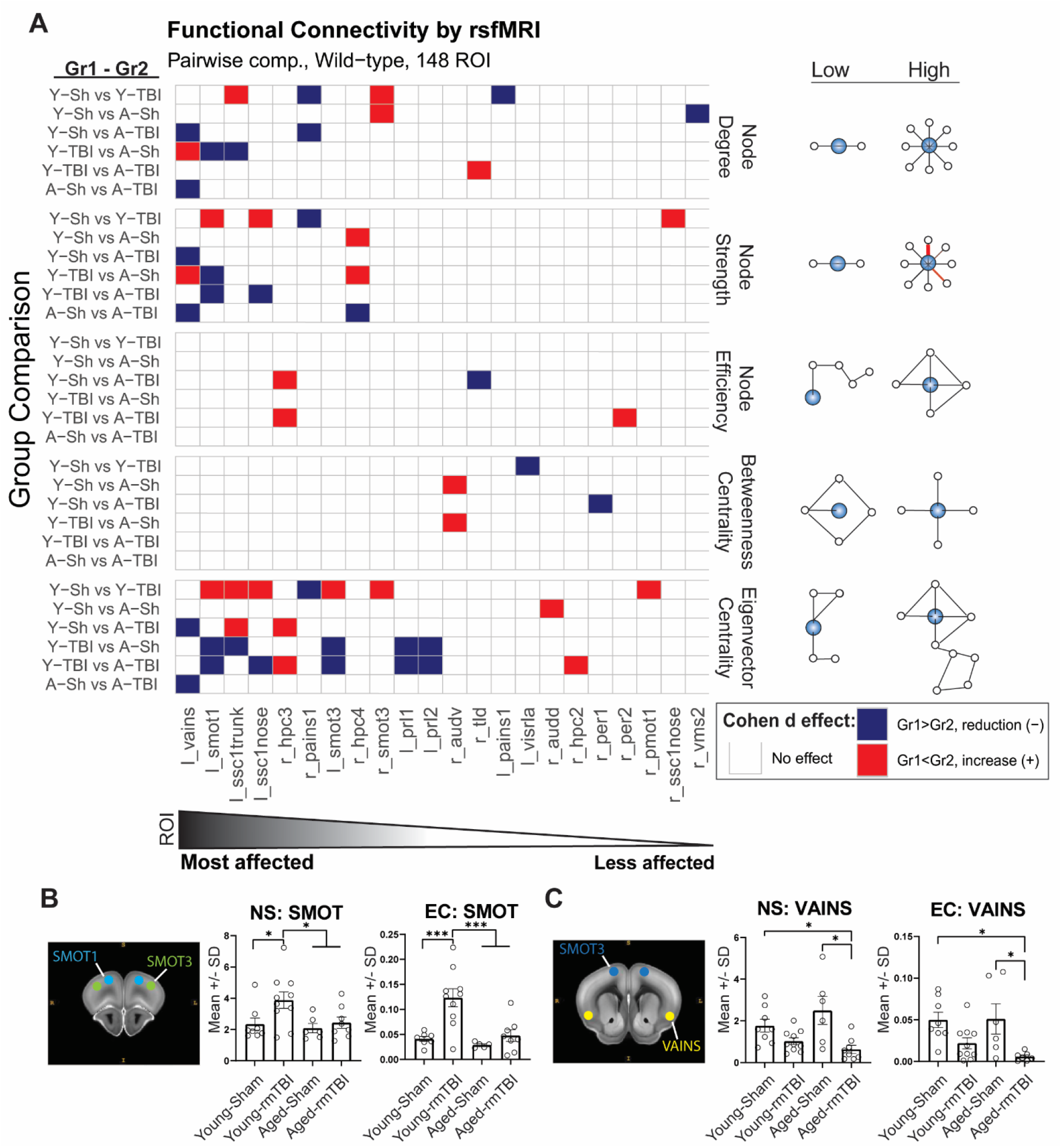
Multiple Comparison and Effect Sizes on Top Affected Brain Regions in Young and Aged Mice After rmTBI. (**A**) Multiple comparisons were conducted on the regions of interest (ROI) showing significant main effects in the two-way ANOVA. Adjusted p-values resulting from pairwise comparisons with Benjamini-Hochberg (FDR-BH) correction are color-coded in the heatplot. The magnitude and direction of the effect were determined by Cohen’s d effect size. Red represents an effect size equal to or greater (red) or lower (blue) than +/−0.8 (large effect). ROIs are arranged from left to right based on the highest number of significant p-values across metrics. Diagrams on the right illustrate scores assigned to a region of interest (ROI, blue circle) based on its connectivity to other regions (white circles), ranging from low to high values. (**B**) In young mice, the secondary motor area (left smot1) was the most affected region by rmTBI, while (**C**) in the aged group, the ventral agranular insula (left vains) took the top spot. NS = node strength, EC = eigenvector centrality. Bar graphs represent the mean with standard deviation (SD). Statistically significant p-values from two-way ANOVA are represented as follows: *p < 0.05, ***p < 0.001. Gr1, group1 is first from left to right in panel; Young-Sham (Y-Sh), n = 7; Young-rmTBI (Y-TBI), n = 10; Aged-Sham (A-Sh), n = 5; Aged-rmTBI (A-TBI), n = 8.

Mean connectivity broadly defines the degree of interconnections between these regions of interest within the brain network, whereas eigenvector centrality describes the quality and influence of these connections (**Figure 4A**). Overall, global mean connectivity and eigenvector centrality revealed age-related differences, where aged mice exhibit denser connectivity in the caudal area while effects in young mice were concentrated in frontal regions. Higher edge densities signify a greater degree of connectivity. Two regions may have the same number of connections (nodes 1 and 2 in **Figure 4B**); however, these regions can significantly differ in terms of information flow and influence within the network, as measured by eigenvector centrality. To gain further insight, we assessed eigenvector centrality by grouping ROIs into four subdivisions: hippocampal formation (HPC), thalamus, striatum, and isocortex (**Figure 4C**). Quantitative analysis reveals a stark contrast in network reconfiguration between young and aged mice following injury (**Figure 4D-4G**). Specifically, the most influential regions concentrated in the isocortex and striatum for young mice and in the hippocampal and thalamic area for aged mice, respectively. These findings raise questions about the efficiency of information flow between the frontal cortex and thalamic regions, prompting us to investigate whether microstructural damage in brain tissue may be the contributing factor.

**Figure 4.**
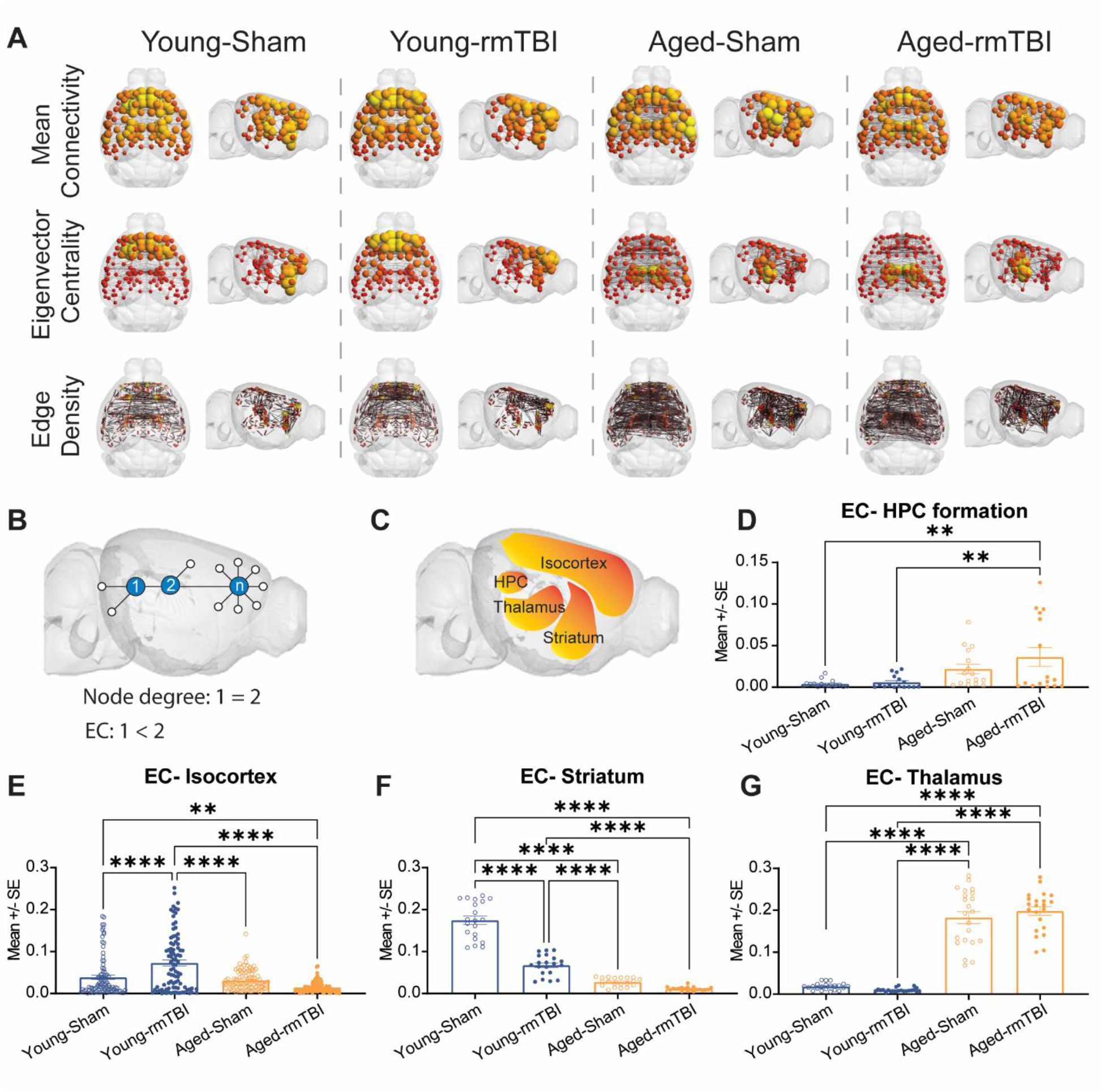
Functional Brain Connectivity by Subdivisions. (**A**). Connectome maps illustrating the average functional connectivity, eigenvector centrality, and total edge densities among all ROIs within the groups. (**B**) A basic diagram depicting node degree and eigenvector centrality relationship. (**C**) Brain areas were further dissected into subdivisions showing age differences in eigenvector centrality in the (**D**) hippocampal formation, (**E**) isocortex, (**F**) striatum, and (**G**) thalamus. Statistically significant p-values from two-way ANOVA are represented as follows: **p < 0.01, ****p < 0.0001. EC = eigenvector centrality, HPC, hippocampus; ROI, regions of interest.

### Age-related disruptions in white matter integrity

Diffusion of water molecules, as an indicator of white matter organization, was examined in 36 regions using DTI (**Figure 5A**). By measuring fractional anisotropy (FA), we inspected diffusion through white matter tracts in a single direction (FA =1, highly organized) or in all directions (FA = 0, tissue damage). Radial and axial diffusivity are sensitive to axonal disruption detecting the flow perpendicular (RD) or parallel (AD) to fiber tracts. Major age-dependent disruptions in diffusion manifested in the corticospinal tract, hippocampal commissure, anterior cingulate, and optic tract. Surprisingly, mean diffusivity (MD), which displays the overall diffusion of water in these regions, was affected uniquely by age (**Figure 5B**). NODDI provided better insight into the density and orientation of axons in intracellular organization (axons and neurites; OD), intracellular fluid (volume occupied by neurites; ICVF) and extracellular (surrounding of neurites; ISOVF) compartments. As in DTI results, age remained as the determinant factor on the neurite density (**Figure 5B**).

**Figure 5.**
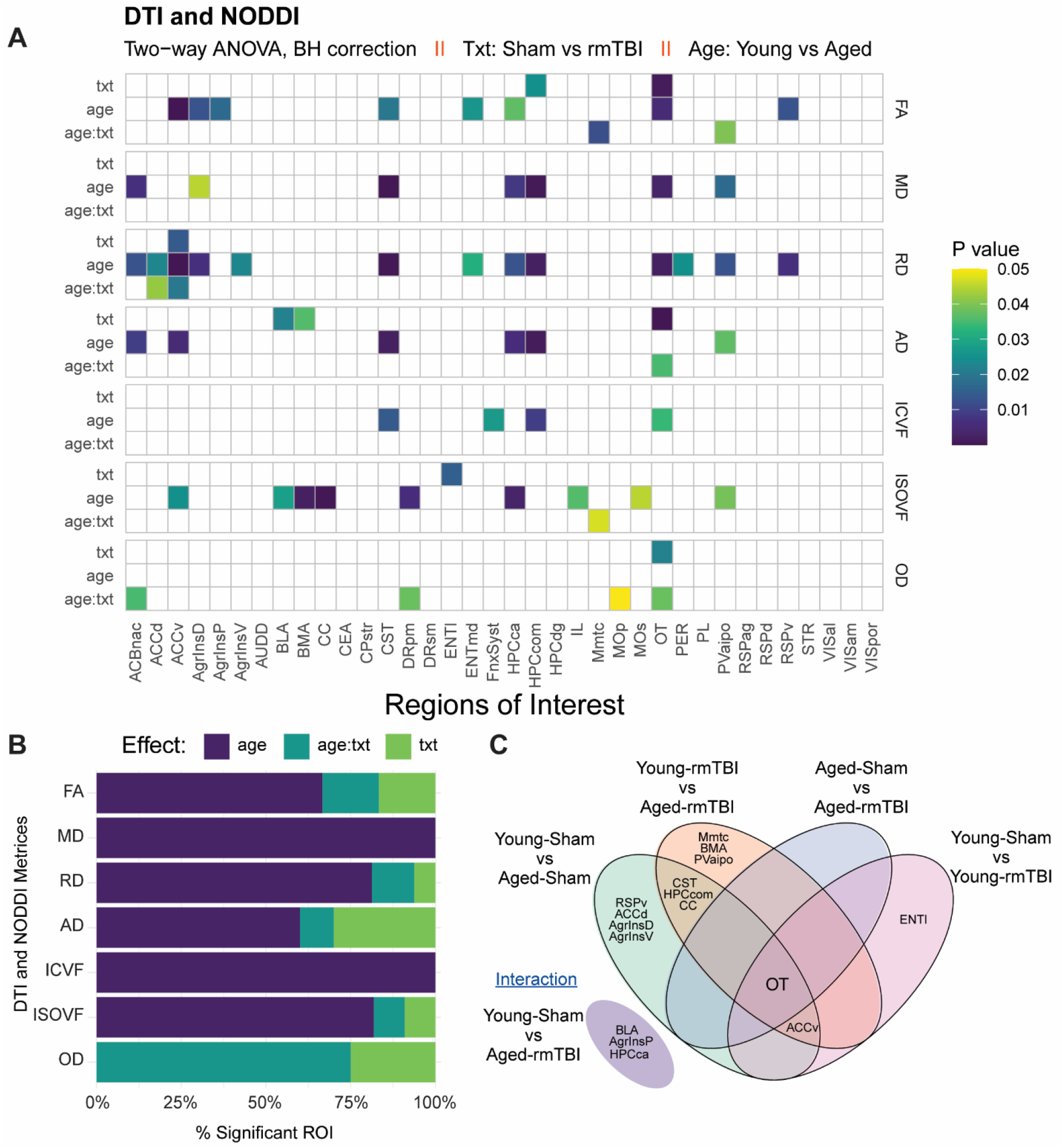
Microstructural Changes in ROIs: DTI and NODDI Metrics. (**A**) Main effects and interactions on each region of interest (ROI)’s microstructure. Significant adjusted p-values resulting from two-way ANOVA (type III) with Benjamini-Hochberg (FDR-BH) correction are color-coded in the heatplot. Diffusion tensor imaging (DTI) metrics include fractional anisotropy (FA, measures directional water diffusion), mean diffusivity (MD, reflects overall water diffusion.), axial diffusivity (AD, measures diffusion along the main voxel axis), and radial diffusivity (RD, analyzes diffusion perpendicular to the main voxel axis). The three compartments measured in neurite orientation dispersion and density imaging (NODDI) analysis were: isotropic volume fraction (ISOVF, quantifies non-neurite components), intracellular volume fraction (ICVF, measures volume taken by neurite), and orientation dispersion index (OD, indicate neurite alignment and organization). (**B**) The percentage of significant ROIs affected by age (young vs aged), txt (sham vs rmTBI) and interaction between these factors. (**C**) Shared regions affected between groups demonstrating the unique effect by age and treatment in white matter integrity. Young-Sham (n = 5), Young-rmTBI (n = 4), Aged-Sham (n = 5), and Aged-rmTBI groups (n = 7). **Supplemental Table 2** describe ROI names in DTI and NODDI analyses.

In control animals, the anterior cingulate, agranular insula, and retrosplenial areas showed a notable decrease in MD and RD due to aging (**Figure 5C**, **Figure 6A**). Conversely, the entorhinal cortex, anterior cingulate, and optic tract were the areas affected in young mice. In line with previous studies, the optic tract was the top affected region showing changes in young and aged animals (**Figure 6A**)^17,35^. Further analysis indicated that aging represents a general increase in fractional anisotropy (FA) and decreased RD when comparing young to aged animals. Independent of injuries, aged animals showed an inverse correlation between FA and RD in secondary motor area, corticospinal tract, and hippocampal commissure (**Figure 6B-6D**). This inverse relationship signifies a noticeable alteration in white matter integrity, distinguishing it from the young group. Interestingly, the interplay between age and injury had a discernible impact on diffusivity in the retrosplenial cortex, a region crucial for diverse processing within the limbic system (**Figure 6E-6F**). Overall, the observed FA/RD relationship is a well-supported indicator of demyelination and axonal damage^35^. Consequently, we conducted additional assessments to explore inflammation as a potential mechanism driving axonal damage.

**Figure 6.**
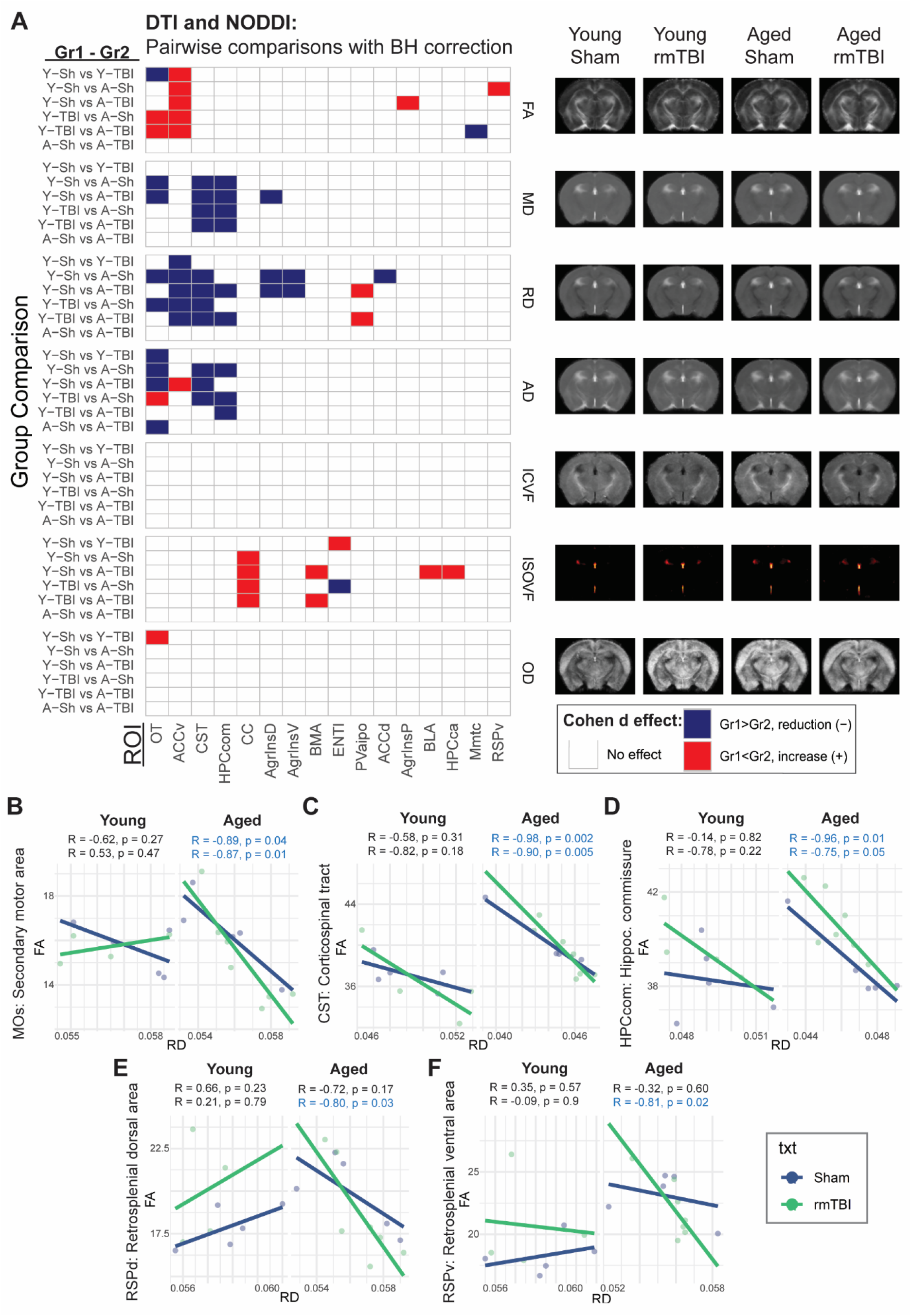
Optic Tract and Anterior Cingulate: Key Regions in DTI and NODDI Analysis Comparison. (**A**) Multiple comparison tests were conducted on the regions of interest (ROI) displaying significant main effects in the two-way ANOVA for both DTI and NODDI analyses. DTI measures include fractional anisotropy (FA, directional water diffusion), mean diffusivity (MD, overall water diffusion), axial diffusivity (AD, diffusion along the main voxel axis), and radial diffusivity (RD, diffusion perpendicular to the main voxel axis). NODDI measures comprise isotropic volume fraction (ISOVF, non-neurite components), intracellular volume fraction (ICVF, volume occupied by neurites), and orientation dispersion index (OD, neurite alignment and organization). Significant effects with a large effect size (Cohen’s d) are indicated in color, while non-effects are shown in white. Adjusted p-values were corrected using Benjamini-Hochberg (FDR) correction. Red signifies an effect size equal to or greater than 0.8, and blue indicates an effect size equal to or less than −0.8. ROIs are ordered based on the highest number of significant p-values, highlighting the optic tract (OT) and anterior cingulate (ACC) as the most affected regions. Representative images correspond to their respective metrics. (**B-D**) Inverse Pearson correlations between FA and RD for the motor area (MOs), corticospinal tract (CST), hippocampal commissure (HPComm) indicates an overall reduction of RD by aging. (**E-F**) Interaction between aging and brain injury augments inverse correlations between FA vs RD signal in retrosplenial area (RSP). Gr1, group1 is first from left to right in panel; Young-Sham (Y-Sh), n =7; Young-rmTBI (Y-TBI), n = 10; Aged-Sham (A-Sh), n = 5; Aged-rmTBI (A-TBI), n = 8.

### Inflammation correlates with higher dispersion in the optic tract

Inflammation is a prevailing pathological response found in aged or injured brains^17,33,36^. Like young mice^17^, injury increased levels of Iba-1 and GFAP inflammatory markers in the optic tract of aged animals (**Figure 7A-7D**). The upregulation of these markers in the optic tract suggests a concerted effort of the immune system to respond to injury-induced stress, implying potential implications for the integrity and function of this neural pathway. For instance, neuroinflammation and neurodegeneration in the optic tract can coincide with damage in other areas like in thalamic nuclei affecting this pathway communication ^37^.

**Fig. 7.**
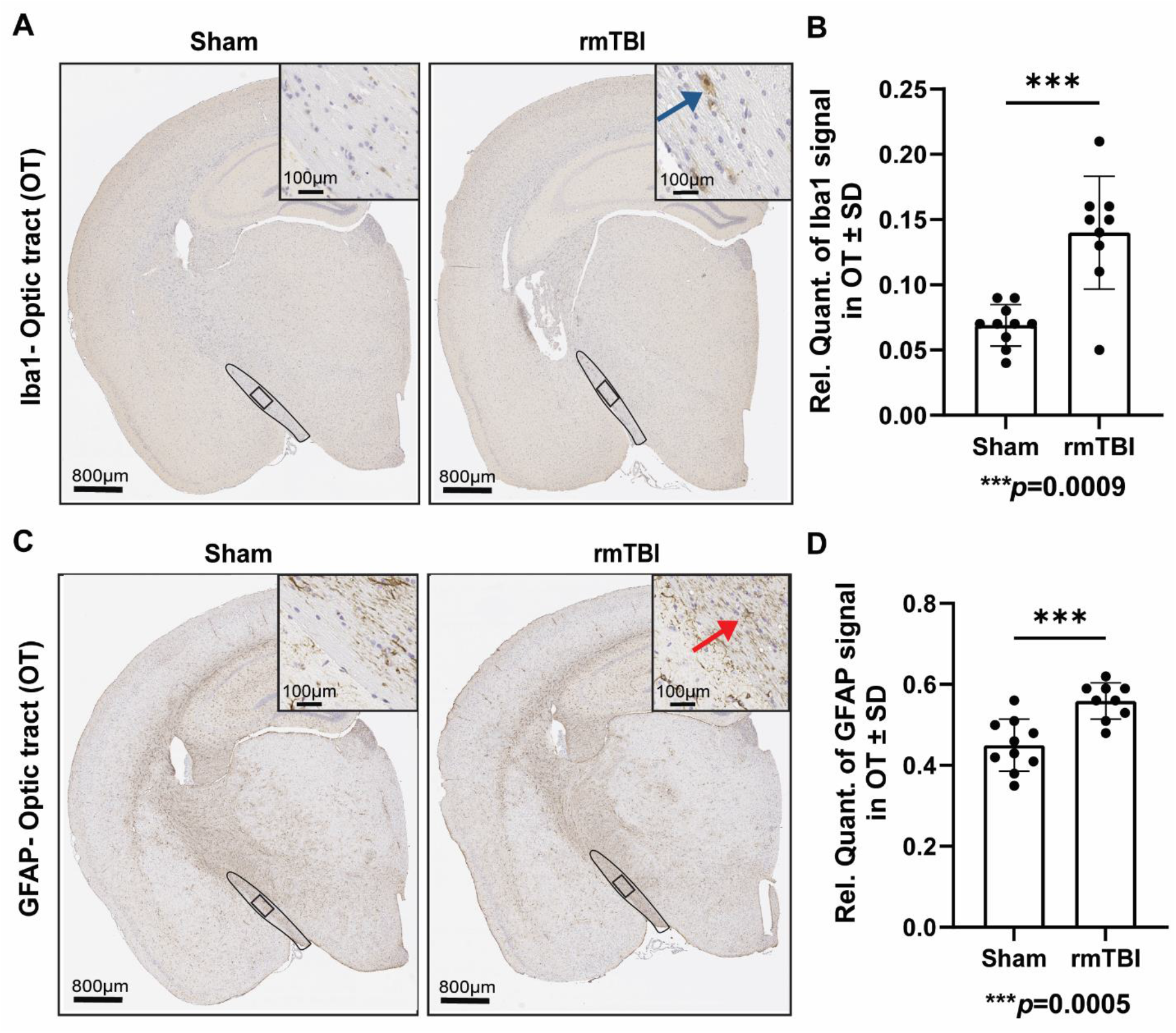
Gliosis is Induced by rmTBI in Optic Tract. (**A**). Following rmTBI, aged mice displayed an increase in (**A-B**) Iba-1 and (**C-D**) GFAP expression in the optic tract. Microglia are indicated by the blue arrowhead, and astrocytes by the red arrowhead. Brain coronal view: 800 μm; Scale bar for inset (enlarged view of the optic tract) = 100 µm. Statistical analysis performed using unpaired t-test with Welch’s correction; Aged-Sham (n = 10), Aged-rmTBI (n =9). SD: standard deviation; Iba-1, Ionized calcium binding adaptor molecule 1; GFAP, Glial fibrillary acidic protein; μm, micrometers; OT, optic tract.

## Discussion

In this study, we investigated alterations in functional connectivity and white matter integrity resulting from rmTBI and aging. We established that major differences in these conditions were not global (**Figure 1**); instead, a more granular evaluation of networks is necessary to define these changes, which in turn underlie the effects of age and head injury on brain health. We established that SWI is reduced by injury (in young animals) and age, disrupting the balance between local clustering and global integration, thereby affecting communication across the entire network. Our observations align with findings in healthy individuals, where global efficiency remains unchanged while changes are detected at local level by decreasing regions efficiency with aging^38^. In the context of mild TBI, it is essential to emphasize that symptoms and visible signs of injuries can manifest days after the event. These effects could intensify and persist for months^19,39^. Undetectable global changes, as revealed in this study, may contribute to why conventional clinical interventions often overlook and fail to accurately diagnose concussions within the first days post-injury. Our findings corroborate in more granular detail that repetitive head injuries alter numerous brain regions leading to reorganization of the entire brain network^17^. More importantly, our results show that brain network restructuring occurs differently between young and aged mice. For instance, young animals primarily exhibited alterations in frontal - motor areas, whereas aged mice showed changes in the agranular insula. A study in healthy young and aged individuals indicated that while global connectivity remains unchanged, the intermodular connections differ by age groups^38^. This local hub reorganization correlates with early signs of cognitive impairments following mTBI^18^. Highlighting contrasting areas affected by rmTBI may predict rmTBI-associated manifestations in behavior and cognition.

The observation of age-related changes in the functioning of the insula is important, given its role in integrating sensory, emotional, cardiovascular and motor functions^40^. The insula is strongly interconnected with the well-established default mode network, often referred to as a “resting state network”^40,41^. Furthermore, aging is linked with declined default-mode network, reduced motor performance, and weakened frontal cortex connections, which are all associated with insular cortex function^40,42–44^. Furthermore, abnormally reduced connections within the anterior cingulate cortex are anticipated with aging. Similar reductions along with decreased segregation and local efficiency are also observed in healthy individuals without injury^38,45^. Additional studies support that aging reduces default-mode network connectivity, motor performance, and frontal cortex connections^42–44^.

It is plausible that the age-related variations we observed in this study are linked to reduced motor plasticity and tissue atrophy in the older cohort, consistent with the effects of aging^33,46,47^. Also, age-related changes in the anterior cingulate cortex impact connectivity between these crucial brain network hubs. Young animals exhibited higher FA and lower RD in ventral anterior cingulate cortex, which is related to improved organization and connectivity in white matter tracts. These changes were absent in aged mice. Two potential explanations are that 1) young animals have a higher degree of plasticity to compensate lost connections following injury and 2) age limits plasticity in the anterior cingulate cortex or motor cortex. Hence, the predominant rmTBI-effects observed in white matter integrity were found in young mice, possibly attributable to pre-existing damage in aged brains. Interestingly, the interplay between aging and injury, rather than each factor individually, significantly influenced tissue diffusion in key limbic regions - the amygdala (BLA), hippocampus (CA area), and posterior insula (**Figure 6A**). These regions play pivotal roles in emotional responses, spatial learning, declarative memory formation, and sensory integration (encompassing awareness of both internal and external bodily states). Our results affirm that the concurrent influence of age and injury leads to a more profound damage in emotional and cognitive processes.

Observations suggest the involvement of thalamic nuclei in conveying age-related connectivity differences. In young mice, the motor cortex and neighboring frontal regions showed increased reorganization. Conversely, aged animals exhibited higher integration (eigenvector centrality) in the thalamic area. Intriguingly, the insular cortex, whose centrality was reduced after rmTBI in aged mice, also shares bidirectional connections with both motor and thalamic areas^48^. The motor cortex and thalamus are not only functionally connected through lifespan, but also play pivotal roles in learning and decision-making^49,50^. Thalamic hyperconnectivity correlates with chronic symptoms in mTBI patients, suggesting that early changes in thalamic nuclei may increase vulnerability to injury^11^. Even without injury, aging impacts the thalamus in healthy subjects by reducing its volume as well as reducing projections to frontal areas, seen during resting or cognitive tasks^42,51^. Consequently, disruption in thalamic-cortical communication can be the underline root for the dissociation in centrality observed in this study.

Other cellular and molecular processes may be implicated in brain connectivity outcomes. For instance, neuroinflammation, oxidative stress, and neuronal loss are common effects related to aging^44^. All these processes are more pronounced in aged brains making them susceptible to damage, dissociation connections, and affecting the capacity to recovery^31,52^. Particularly, inflammation in the optic tract likely affects the visual and sensory input integration. This is noteworthy because reduced visual acuity, altered visual processing and other eye conditions are more prevalent in elderly individuals. Also, higher inflammation in optic tract might underlie visual impairment after injury and other symptoms like insomnia, attention deficits, and memory problems. Enhanced connectivity in this area could serve as a compensatory response to inflammation or a temporary adjustment during the initial days after injury. This could explain the mild symptoms in vision during early days as the brain adjusts.

### Future studies and limitations

Whether the centrality observed in the thalamus of aged mice results from damaged connections remain uncertain. Ongoing studies to investigate whether centrality represents a compensatory mechanism or delayed communication between thalamic-frontal connections are underway. Further insights can be gained by assessing different thalamic and main connecting tracts, like corpus callosum, to pinpoint specific regions responsible for the dissociation between frontal and thalamic connectivity. Notably, there has been a greater number of longitudinal studies conducted in young rodents compared to aged mice. The present study is one of limited investigations assessing changes at 22 months of age. This presents challenges in predicting and comparing long-term effects in this age group. Also, the onset of strain-related diseases occurring beyond this timepoint can be misleading of the true impact of injury. Discussions regarding the lasting effects by rmTBI have been postulated in recent years. This is because others using closed head injuries detect recovery after a month following the injury event^53,54^. However, here we use the surgery-free CHIMERA model which produces lasting effects in tissue pathology, imaging, neurophysiology, and behavior^55–57^. Lastly, to enhance the translational value of our findings, it would be advantageous to link them with other brain and blood markers.

## Conclusion

In summary, our study reveals age-specific changes in brain connectivity and white matter properties following multiple mild traumatic brain injuries. Using a multi-modal approach, we identify reduced communication efficiency and contrasting effects on the small-world index, influencing network segregation with age. These discoveries shed light on the intricate interactions between age, injuries, and brain connectivity, offering implications for understanding the solid effects of rmTBIs in the context of aging and the potential early pathological features that may develop to neurodegeneration and contribute to impairment of brain functions.

### Data Access Statement

We are dedicated to the broad and open dissemination of our research outcomes. In line with this, we will adhere to the NIH Grant Policy on Sharing of Unique Research Resources, which involves the Principles and Guidelines for Recipients of NIH Research Grants and Contracts on Obtaining and Disseminating Biomedical Research Resources (1999). Our data is securely stored in our institutional network computers and HiPerGator supercomputer. Access to data will be made possible after publication, in agreement with the policies established by the University of Florida. The corresponding author will be the contact to access and grant permission to transfer files. If any intellectual property necessitates a patent, we will ensure that the technology, materials, and data are accessible to the broader research community in accordance with the NIH Principles and Guidelines document.

### Transparency, Rigor, and Reproducibility

For imaging studies, the total sample size was 33 consisting of Young-Sham (n = 8), Young-rmTBI (n = 10), Aged-Sham (n = 7), and Aged-rmTBI (n = 8). For IHC experiment in aged mice, the total sample size was 19 (Sham, n = 10; rmTBI, n = 9). Animals were randomly assigned to experimental groups, and the injury procedures and imaging processing were designed and conducted with blinding. Information about the animals’ group assignments was not revealed until the statistical analyses were conducted.

## Acknowledgements

We thank Dave Borchelt and Susan Fromholt for their invaluable contribution in supplying the aged animals, crucial for the successful completion of this study. The main funding for this project was provided by NIH/NIA grant 1 R01 AG074584-01 and partial support from the NIH grant S10 RR025671 for MRI/S instrumentation. We extend our gratitude to the Alzheimer’s Association for the 2019-AARFD-644407 fellowship grant, awarded to Marangelie Criado-Marrero, which greatly supported her training and the research findings detailed in this publication. A portion of the experiments were conducted at the McKnight Brain Institute, within the Advanced Magnetic Resonance Imaging and Spectroscopy (AMRIS) Facility at the National High Magnetic Field Laboratory. This facility was made possible through additional support from the National Science Foundation Cooperative Agreement DMR-1644779 and the State of Florida.

## Author contributions

Conceptualization of this study was led by MCM, SR, and JFA. Experiments were performed by SR and DB. Imaging data was analyzed by MCM and EB. Manuscript was prepared by MCM and JFA. CW previously provided the CHIMERA device. All authors contributed to final editing of the current manuscript and approved the submitted version.

