## Supplemental Information for "Age dictates brain functional connectivity and axonal integrity following repetitive mild traumatic brain injuries"

**Supplementary information:**

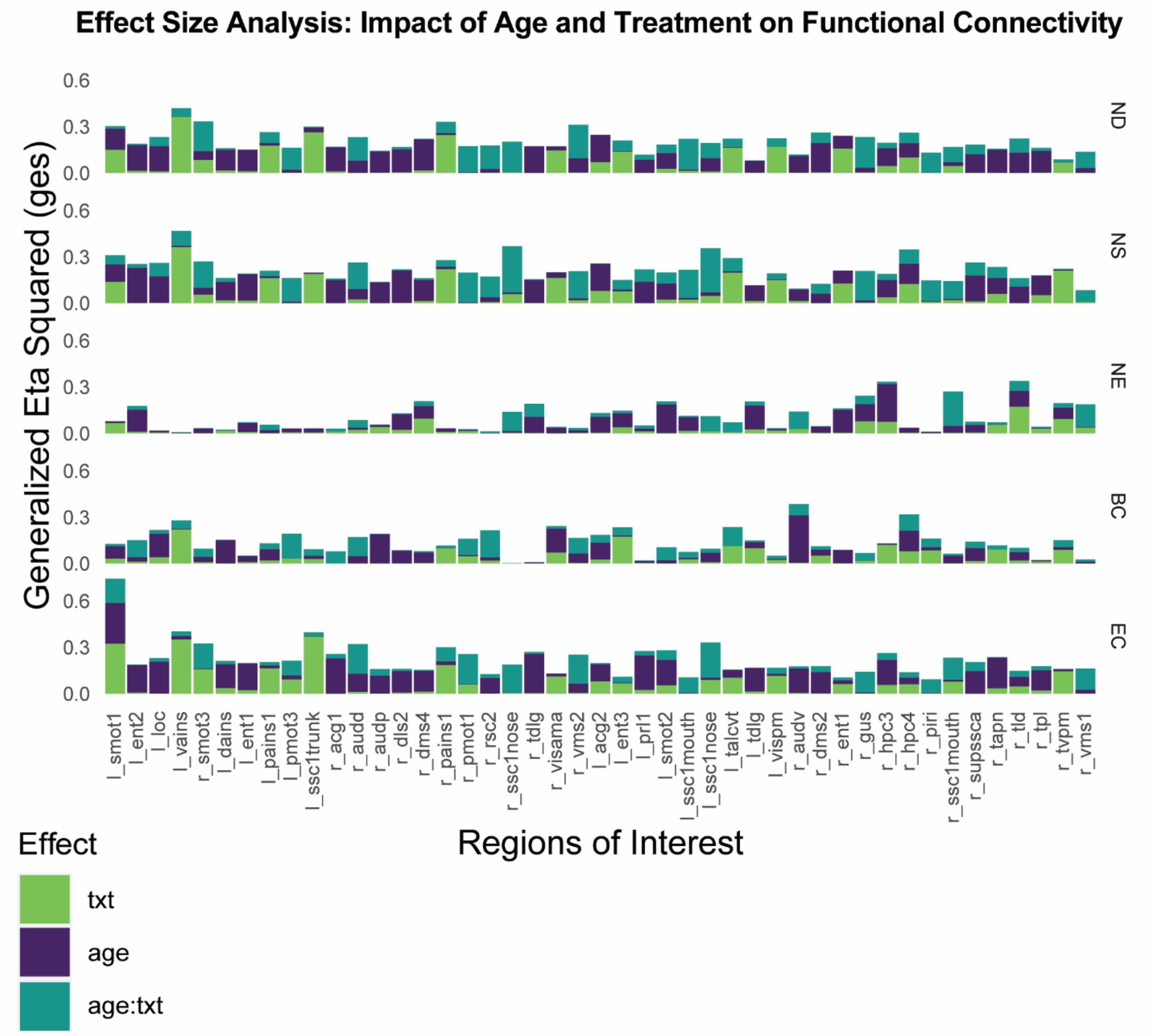


**Figure S1.** Generalized eta squared distributions for main effects on each ROI measured through functional connectivity. The metrices for rsfMRI include: node degree (ND), node strength (NS), node efficiency (NE), betweenness centrality (BC), and eigenvector centrality (EC).

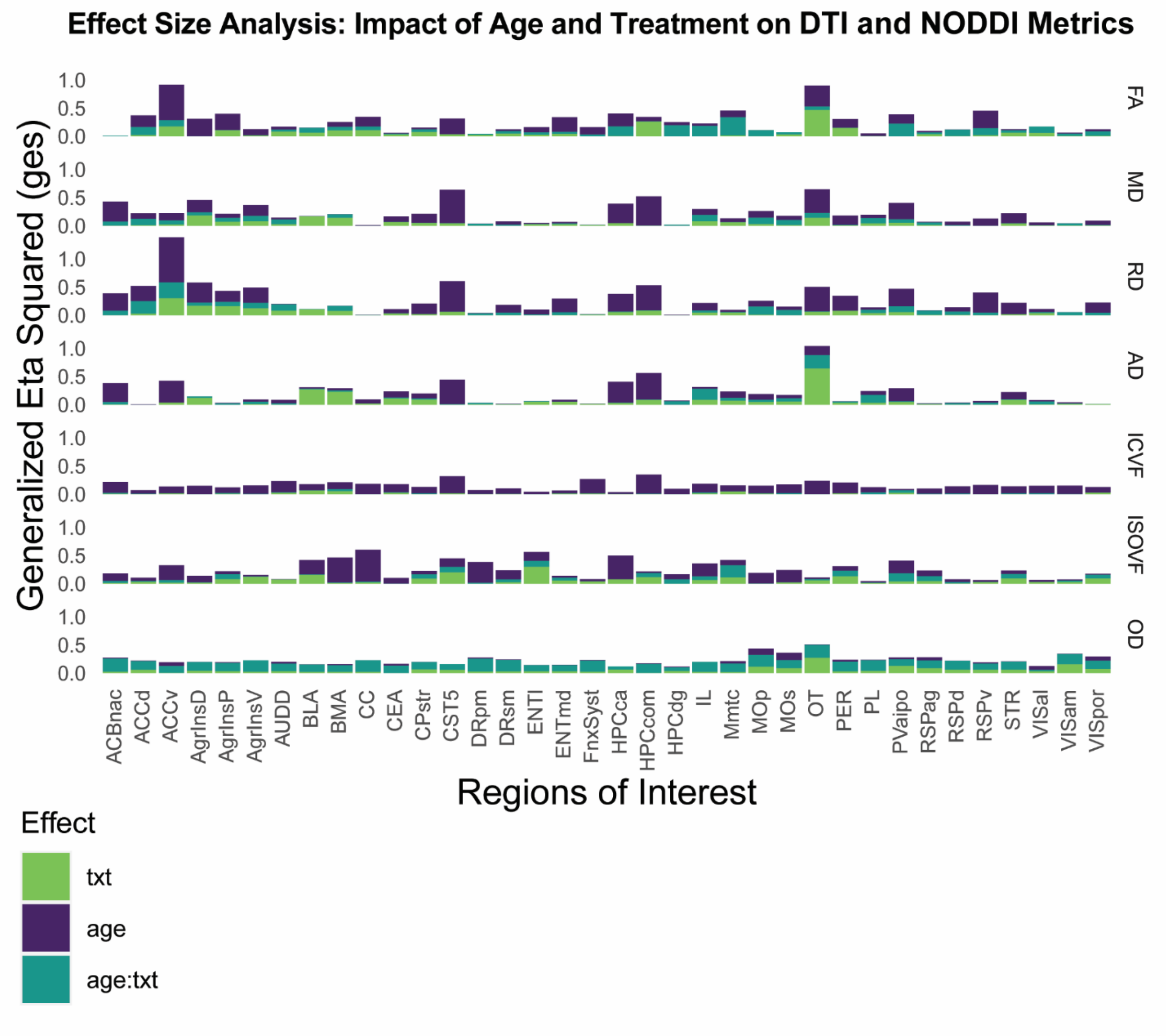


**Figure. S2.** Generalized eta squared distributions for main effects on each ROI evaluated through DTI and NODDI analyses. Fractional anisotropy (FA), mean diffusivity (MD), axial diffusivity (AD), and radial diffusivity (RD). The three compartments measured in neurite orientation dispersion and density imaging (NODDI) analysis were: isotropic volume fraction (ISOVF), intracellular volume fraction (ICVF), and orientation dispersion index (OD).

__Table legend__

**Supplemental Table 1. Regions of interests used in rsfMRI.** A comprehensive list of 148 regions (ROI) located in left (L) and right (R) hemispheres were analyzed using rsfMRI and graph theory.

| Table S1: Regions of interest (ROI) in rsfMRI analysis | | | |
|---|---|---|---|
| **Order** | **Hemisphere** | **ROI (sphere location)** | **Abbreviation** |
| 1 | Right | Hippocampus | r_hpc1 |
| 2 | Right | Hippocampus | r_hpc2 |
| 3 | Right | Hippocampus | r_hpc3 |
| 4 | Right | Hippocampus | r_hpc4 |
| 5 | Right | Hippocampus | r_hpc5 |
| 6 | Right | Entorhinal | r_ent1 |
| 7 | Right | Entorhinal | r_ent2 |
| 8 | Right | Entorhinal | r_ent2 |
| 9 | Right | Perirhinal | r_per1 |
| 10 | Right | Perirhinal | r_per2 |
| 11 | Right | Temporal | r_temp |
| 12 | Right | Retrosplenial | r_rsc1 |
| 13 | Right | Retrosplenial | r_rsc2 |
| 14 | Right | Retrosplenial | r_rsc3 |
| 15 | Right | Anterior Cingulate | r_acg1 |
| 16 | Right | Anterior Cingulate | r_acg2 |
| 17 | Right | Prelimbic | r_prl1 |
| 18 | Right | Prelimbic | r_prl2 |
| 19 | Right | Infralimbic | r_il |
| 20 | Right | Lateral Orbital | r_loc |
| 21 | Right | Ventrolateral Orbital | r_voc |
| 22 | Right | Dorsal Agranular Insula | r_dains |
| 23 | Right | Posterior Agranular Insula | r_pains1 |
| 24 | Right | Posterior Agranular Insula | r_pains2 |
| 25 | Right | Ventral Agranular Insula | r_vains |
| 26 | Right | Visual Anterior area | r_visaa |
| 27 | Right | Visual Rostrolateral area | r_visrla |
| 28 | Right | Visual Anteromedial area | r_visama |
| 29 | Right | Visual Lateral area | r_visla |
| 30 | Right | Visual primary visual | r_vispv1 |
| 31 | Right | Visual primary visual | r_vispv2 |
| 32 | Right | Visual Posterolateral | r_vispl |
| 33 | Right | Visual Posteromedial | r_vispm |
| 34 | Right | Visual postrhinal | r_vispr |
| 35 | Right | Auditory dorsal | r_audd |
| 36 | Right | Auditory primary | r_audp |
| 37 | Right | Auditory ventral | r_audv |
| 38 | Right | SSC1 nose | r_ssc1nose |
| 39 | Right | SSC1 barrel | r_ssc1barrel |
| 40 | Right | SSC1 lower limb | r_ssc1lowlimb |
| 41 | Right | SSC1 mouth layer | r_ssc1mouth |

| | | | |
|---|---|---|---|
| 42 | Right | SSC1 upper limb | r_ssc1ulimb |
| 43 | Right | SSC1 trunk | r_ssc1trunk |
| 44 | Right | Supplemental SSC area | r_supssca |
| 45 | Right | Gustatory cortex | r_gus |
| 46 | Right | Visceral area | r_visc |
| 47 | Right | Piriform | r_piri |
| 48 | Right | Motor primary | r_pmot1 |
| 49 | Right | Motor primary | r_pmot2 |
| 50 | Right | Motor primary | r_pmot3 |
| 51 | Right | Motor secondary | r_smot1 |
| 52 | Right | Motor secondary | r_smot2 |
| 53 | Right | Motor secondary | r_smot3 |
| 54 | Right | Thalamus mediodorsal | r_tmd |
| 55 | Right | Thalamus anterolateral complex (ventral) | r_talcvt |
| 56 | Right | Thalamus laterodorsal | r_tld |
| 57 | Right | Thalamus posterior complex | r_tpc |
| 58 | Right | Thalamus ventral posteromedial | r_tvpm |
| 59 | Right | Thalamus paracentral nucleus | r_tpcn |
| 60 | Right | Thalamus posterior lateral | r_tpl |
| 61 | Right | Thalamus central lateral – parafascicular | r_tclp |
| 62 | Right | Thalamus anterior pretectal nucleus | r_tapn |
| 63 | Right | Thalamus dorsal lateral geniculate | r_tdlg |
| 64 | Right | Thalamus ventromedial | r_tvm |
| 65 | Right | Dorsomedial Striatum | r_dms1 |
| 66 | Right | Dorsomedial Striatum | r_dms2 |
| 67 | Right | Ventromedial Striatum | r_vms1 |
| 68 | Right | Dorsolateral Striatum | r_dls1 |
| 69 | Right | Dorsolateral Striatum | r_dls2 |
| 70 | Right | Ventromedial Striatum | r_vms2 |
| 71 | Right | Dorsomedial Striatum | r_dms3 |
| 72 | Right | Dorsomedial Striatum | r_dms4 |
| 73 | Right | Dorsolateral Striatum | r_dls3 |
| 74 | Right | Dorsolateral Striatum | r_dls4 |
| 75 | Left | Hippocampus | l_hpc1 |
| 76 | Left | Hippocampus | l_hpc2 |
| 77 | Left | Hippocampus | l_hpc3 |
| 78 | Left | Hippocampus | l_hpc4 |
| 79 | Left | Hippocampus | l_hpc5 |
| 80 | Left | Entorhinal | l_ent1 |
| 81 | Left | Entorhinal | l_ent2 |
| 82 | Left | Entorhinal | l_ent3 |
| 83 | Left | Perirhinal | l_per1 |
| 84 | Left | Perirhinal | l_per2 |
| 85 | Left | Temporal | l_temp |
| 86 | Left | Retrosplenial | l_rsc1 |

| | | | |
|---|---|---|---|
| 87 | Left | Retrosplenial | l_rsc2 |
| 88 | Left | Retrosplenial | l_rsc3 |
| 89 | Left | Anterior Cingulate | l_acg1 |
| 90 | Left | Anterior Cingulate | l_acg2 |
| 91 | Left | Prelimbic | l_prl1 |
| 92 | Left | Prelimbic | l_prl2 |
| 93 | Left | Infralimbic | l_il |
| 94 | Left | Lateral Orbital | l_loc |
| 95 | Left | Ventrolateral Orbital | l_voc |
| 96 | Left | Dorsal Agranular Insula | l_dains |
| 97 | Left | Posterior Agranular Insula | l_pains1 |
| 98 | Left | Posterior Agranular Insula | l_pains2 |
| 99 | Left | Ventral Agranular Insula | l_vains |
| 100 | Left | Visual Anterior area | l_visaa |
| 101 | Left | Visual Rostrolateral area | l_visrla |
| 102 | Left | Visual Anteromedial area | l_visama |
| 103 | Left | Visual Lateral area | l_visla |
| 104 | Left | Visual primary visual | l_vispv1 |
| 105 | Left | Visual primary visual | l_vispv2 |
| 106 | Left | Visual Posterolateral | l_vispl |
| 107 | Left | Visual Posteromedial | l_vispm |
| 108 | Left | Visual postrhinal | l_vispr |
| 109 | Left | Auditory dorsal | l_audd |
| 110 | Left | Auditory primary | l_audp |
| 111 | Left | Auditory ventral | l_audv |
| 112 | Left | SSC1 nose | l_ssc1nose |
| 113 | Left | SSC1 barrel | l_ssc1barrel |
| 114 | Left | SSC1 lower limb | l_ssc1lowlimb |
| 115 | Left | SSC1 mouth layer | l_ssc1mouth |
| 116 | Left | SSC1 upper limb | l_ssc1ulimb |
| 117 | Left | SSC1 trunk | l_ssc1trunk |
| 118 | Left | Supplemental SSC area | l_supssca |
| 119 | Left | Gustatory cortex | l_gus |
| 120 | Left | Visceral area | l_visc |
| 121 | Left | Piriform | l_piri |
| 122 | Left | Motor primary | l_pmot1 |
| 123 | Left | Motor primary | l_pmot2 |
| 124 | Left | Motor primary | l_pmot3 |
| 125 | Left | Motor secondary | l_smot1 |
| 126 | Left | Motor secondary | l_smot2 |
| 127 | Left | Motor secondary | l_smot3 |
| 128 | Left | Thalamus mediodorsal | l_tmd |
| 129 | Left | Thalamus anterolateral complex (ventral) | l_talcvt |
| 130 | Left | Thalamus laterodorsal | l_tld |
| 131 | Left | Thalamus posterior complex | l_tpc |

| | | | |
|---|---|---|---|
| 132 | Left | Thalamus ventral posteromedial | l_tvpm |
| 133 | Left | Thalamus paracentral nucleus | l_tpcn |
| 134 | Left | Thalamus posterior lateral | l_tpl |
| 135 | Left | Thalamus central lateral – parafascicular | l_tclp |
| 136 | Left | Thalamus anterior pretectal nucleus | l_tapn |
| 137 | Left | Thalamus dorsal lateral geniculate | l_tdlg |
| 138 | Left | Thalamus ventromedial | l_tvm |
| 139 | Left | Dorsomedial Striatum | l_dms1 |
| 140 | Left | Dorsomedial Striatum | l_dms2 |
| 141 | Left | Ventromedial Striatum | l_vms1 |
| 142 | Left | Dorsolateral Striatum | l_dls1 |
| 143 | Left | Dorsolateral Striatum | l_dls2 |
| 144 | Left | Ventromedial Striatum | l_vms2 |
| 145 | Left | Dorsomedial Striatum | l_dms3 |
| 146 | Left | Dorsomedial Striatum | l_dms4 |
| 147 | Left | Dorsolateral Striatum | l_dls3 |
| 148 | Left | Dorsolateral Striatum | l_dls4 |

**Supplemental Table 2. Regions of interests examined DTI and NODDI analyses.** A total of 36 regions of interest (ROI) including white and grey matter areas were examined.

| **Table S2: Regions of interest (ROI) in DTI and NODDI analysis** | | | |
|---|---|---|---|
| **Order** | | **ROI** | **Abbreviation** |
| 1 | | Anterior cingulate area- dorsal part | ACCd |
| 2 | | Anterior cingulate area- ventral part | ACCv |
| 3 | | Agranular insular area- dorsal part | AgrInsD |
| 4 | | Agranular insular area- posterior part | AgrInsP |
| 5 | | Hippocampus | AgrInsV |
| 6 | | Dorsal auditory area | AUDD |
| 7 | | Basolateral amygdala | BLA |
| 8 | | Basomedial amygdala | BMA |
| 9 | | Corpus collosum | CC |
| 10 | | Central medial amygdala | CEA |
| 11 | | Entorhinal cortex- lateral layer | ENTl |
| 12 | | Entorhinal cortex- medial dorsal layer | ENTmd |
| 13 | | Fornix system | FnxSyst |
| 14 | | Hippocampus – CA1 region | HPCca |
| 15 | | Hippocampus – dentate gyrus | HPCcdg |
| 16 | | Hippocampal commissure | HPCcomm |
| 17 | | Infralimbic cortex | IL |
| 18 | | Mammillothalamic tract | Mmtc |
| 19 | | Primary motor area | MOp |
| 20 | | Secondary motor area | MOs |

| | | | |
|---|---|---|---|
| 21 | | Perirhinal area | PER |
| 22 | | Prelimbic cortex | PL |
| 23 | | Periventricular hypothalamic nucleus | PVaipo |
| 24 | | Retrosplenial area lateral agranular layer | RSPag |
| 25 | | Retrosplenial dorsal area | RSPd |
| 26 | | Retrosplenial ventral area | RSPv |
| 27 | | Thalamus, sensory-motor cortex related | DRsm |
| 28 | | Thalamus, polymodal association cortex related | DRspm |
| 29 | | Anterolateral visual area | VISal |
| 30 | | Anteromedial visual area | VISam |
| 31 | | Postrhinal area | VISpor |
| 32 | | Corticospinal tract | CST |
| 33 | | Nuclues accumbens | ACBnac |
| 34 | | Caudoputamen | CPstr |
| 35 | | Striatum | STR |
| 36 | | Optic Tract | OT |
